# CandiHap: a toolkit for haplotype analysis for sequence of samples and fast identification of candidate causal gene(s) in genome-wide association study

**DOI:** 10.1101/2020.02.27.967539

**Authors:** Xukai Li, Zhiyong Shi, Qianru Qie, Jianhua Gao, Xingchun Wang, Yuanhuai Han

**Affiliations:** College of Life Sciences, Shanxi Agricultural University, Taigu, 030801, China; College of Agriculture, Shanxi Agricultural University, Taiyuan, 030801, China

## Abstract

Genome-wide association study (GWAS) is widely used to identify genes involved in plants, animals and human complex traits. Generally, the identified SNP is not necessarily the causal variant, but it is rather in linkage disequilibrium (LD). One key challenge for GWAS results interpretation is to rapidly identify causal genes and provide profound evidence on how they affect the trait. Researches want to identify candidate causal variants from the most significant SNPs of GWAS in any species and on their local computer, while to complete these tasks are to be time-consuming, laborious and prone to errors and omission. To our knowledge, so far there is no tool available to solve the challenge for GWAS data very quickly. Based on the standard VCF (variant call format) format, CandiHap is developed to fast preselection candidate causal SNPs and gene(s) from GWAS by integrating LD result, SNP annotation, haplotype analysis and traits statistics of haplotypes. Investigators can specify genes or linkage regions based on GWAS results, linkage disequilibrium (LD), and predicted candidate causal gene(s). It supported Windows, Mac and Linux computers and servers in graphical interface and command line, and applied to any other plant, animal or bacteria species. The source code of CandiHap tool is freely available at https://github.com/xukaili/CandiHap

## Introduction

With next generation sequencing (NGS), genome sequencing is becoming inexpensive and routine, and the obtention of large numbers of SNPs is convenient. Genome-wide association study (GWAS) has become established in medical, biological and agricultural research to elucidate the genetic basis of phenotypic traits such as disease or economically important features (Visscher et al. 2012; Visscher et al. 2017). SNP can alter a protein directly (non-synonymous SNPs, stop gained or stop lost SNPs, frameshift SNPs or SNPs in splice sites) or it can implicate gene expression if SNP are located in regulatory regions. From a huge number of genome-wide variants, a GWAS investigation generally identifies a few SNPs that are statistically significantly associated with some trait. As GWAS serves as initializations of future genetic and mechanism study of complex traits, one of the key challenges of GWAS data interpretation is to identify causal SNPs (the SNPs that affect trait) and provide profound evidence and hypothesis on the mechanism through which they affect the trait (McCarthy and Hirschhorn 2008).

There are some researches focusing on inferring candidate causal SNPs from the most significant (SNPs with *P* value below certain threshold) (Hindorff et al. 2009; Li et al. 2012) and prioritizing the most significant SNPs by linkage disequilibrium (LD) analysis and functional SNP annotation (Adzhubei et al. 2010; Johnson et al. 2008; Kumar et al. 2009; Lee and Shatkay 2008; Mi et al. 2010; Saccone et al. 2010; Schmitt et al. 2010; Xu and Taylor 2009; Yuan et al. 2006; Yue et al. 2006). However, most existing tools are web-based tools or command-line for human studies, severely limiting those widely use. In fact, more researches want to identify candidate causal variants affect traits from the most significant SNP of GWAS in their own species (not only for human, but also for any species) and on their local computer (for maintain secrecy), while to complete these tasks are to be time-consuming, laborious and prone to errors and omission and not a convivial interface. A feasible proposal to address the above problems is to development a software to fast find candidate causal variants and gene(s). So far, there is no tool available to provide a solution for GWAS data very quickly. CandiHap aims to provide an open source to facilitate researchers to identify the candidate causal SNPs and gene(s) of traits and to guide future genetic and mechanism study.

## Methods

### Variant filtering and annotation

The variants of VCF file was further filtered using the VCFtools (Danecek et al. 2011) (ver. 0.1.15). The SNPs and indels were considered valid for the study if they met the following requirements: (1) two alleles only; (2) exclude sites on the basis of the proportion of missing data >0.9 (defined to be between 0 and 1, where 0 allows sites that are completely missing and 1 indicates no missing data allowed); (3) minor allele frequency ≥0.05; and mean depth values ≥5. SNPs that did not meet these four criteria were excluded from the study. All identified SNPs that passed quality screening were further annotated with ANNOVAR (ver. 2015 Dec 14) based on the gene annotation of the reference genome (Wang et al. 2010). In practical application, users can adjust the above parameters for a study. When a VCF file is submitted, ANNOVAR is computed to rapidly categorize the effects of variants in the reference genome sequence. ANNOVAR annotates variants based on their genomic locations (annotated genomic locations can be intronic, exonic or intergenic) and predicts coding effects (mainly synonymous or non-synonymous amino-acid replacement). The process can be applied to any other plant, animal or bacteria species, by providing the genome file and its GFF (generic feature format) annotation file.

### Software development

CandiHap is written in Perl 5 (v 5.26, https://www.perl.org), R (v 3.5, https://www.r-project.org) and Python 2.7 (https://www.python.org), which supported Windows, Mac and Linux computers and servers in graphical interface and command lines. Graphics are created by R. The graphical user interface is written in electron, which is freely available and registration is not required. Besides the graphical interface software, users can run CandiHap through command lines by using the Linux or Mac. For a given SNP that was found significant in a GWAS, runtime is ∼1 min for a set of 400 samples and ∼3 million SNPs. The CandiHap tool is freely available at https://github.com/xukaili/CandiHap.

### General statistics

Using perl and R, the results provides and displays various statistics about the haplotypes such as annotation statistics, type of variations, number of varieties, varieties ID and its phenotype, average and SD (standard deviation) of phenotype. A boxplot of gene showed significant difference in the phenotype of each haplotype.

Methods of the graphical interface

## Results

### Process overview

Here, we propose the CandiHap local software, which supported Windows, Mac and Linux computers and servers in graphical interface and command lines. An overview of the process is presented Figure 1. Starting from a VCF file as entry point, the process first annotates the variants using an annotated reference genome to produce a new VCF file from which variants and genotyping data can be then mined and sent into a series of modules in charge of various processes. User has then the possibility to analyze variants either at the genome level or at the gene level. The GWAS result of genomic regions (Fig. 1a) and linkage disequilibrium (LD) can be defined by entering the limits, the application will loop and process these region genes.

**Fig. 1.**
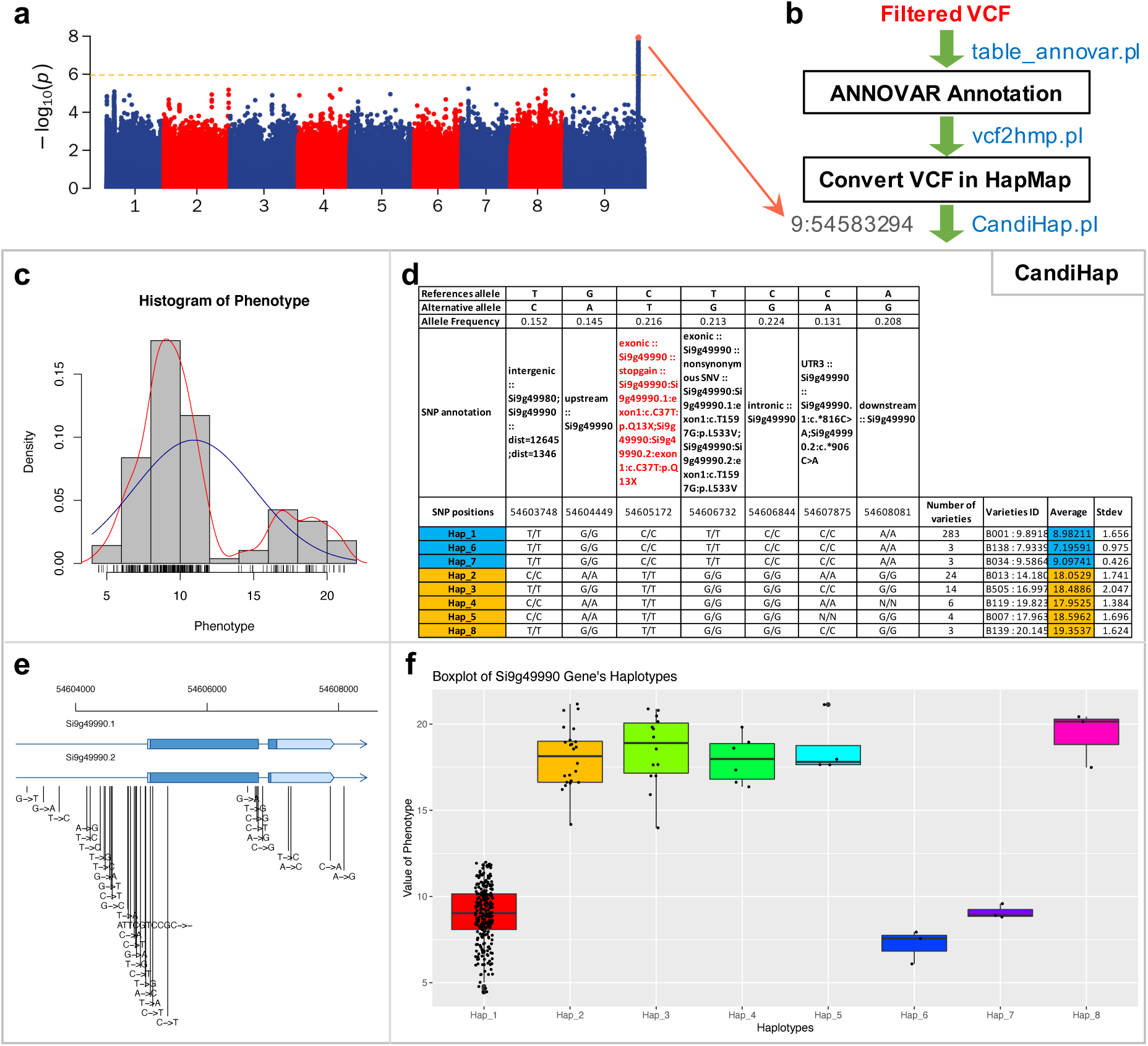
Overview of the CandiHap process. (a) A GWAS result. (b) General schema of the process. (c) The histogram of phenotype. (d) The statistics of haplotypes. (e) Gene structure and SNPs of key gene. (f) Boxplot of key gene’s haplotypes.

**Fig. 1.**
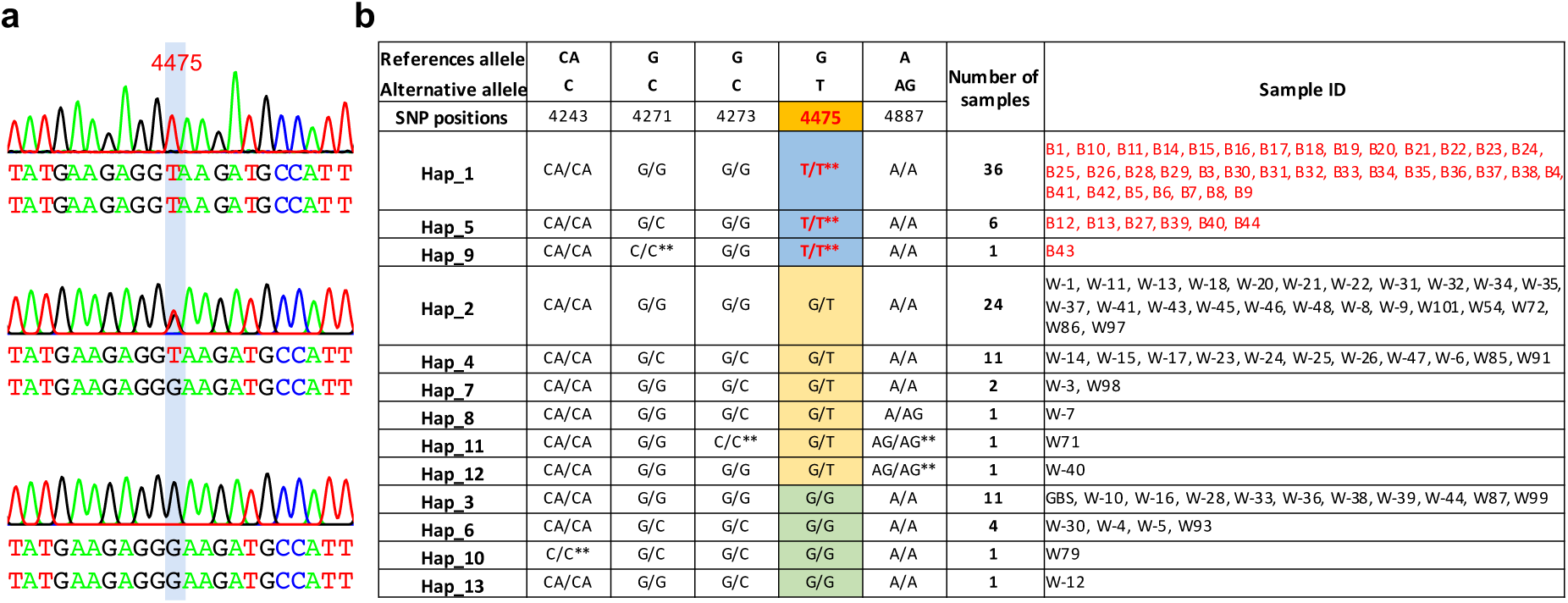
The haplotype analysis in Sanger ab1 files. (a) PeakTrace of ab1 images of three genotypes. (b) The statistics of haplotypes.

### Input data and running procedure

The CandiHap implements a three-stage analysis (Fig. 1b). The first stage is to annotate the VCF file for GWAS by ANNOVAR (table_annovar.pl). The second stage is to convert the txt result of annovar to hapmap format (vcf2hmp.pl). The third stage required input data of hapmap file, GFF file of your reference genome, the phenotype data, the linkage disequilibrium (LD), and a most significant SNPs position of GWAS result. If users only want to run one gene, they can input the vcf, phenotype, gff and gene ID. Besides the graphical interface software, users can run CandiHap through command lines by using the Linux, Mac and DOS.

There are mainly three steps included in the CandiHap analytical through command lines, and the test data files can freely download at https://github.com/xukaili/CandiHap.

1. To annotate the vcf by ANNOVAR: 1.1 gffread test.gff -T -o test.gtf 1.2 gtfToGenePred -genePredExt test.gtf si_refGene.txt 1.3 retrieve_seq_from_fasta.pl --format refGene --seqfile genome.fa si_refGene.txt --outfile si_refGeneMrna.fa 1.4 table_annovar.pl test.vcf ./ --vcfinput --outfile test --buildver si --protocol refGene --operation g-remove
2. To convert the txt result of annovar to hapmap format (0.1 means the minor allele frequency (MAF)): perl vcf2hmp.pl test.vcf test.si_multianno.txt 0.1
3. To run CandiHap: perl GWAS_LD2haplotypes.pl ./test.gff ./haplotypes.hmp ./Phenotype.txt 50kb 9:54583294 Or to run CandiHap by one gene: perl CandiHap.pl ./haplotypes.hmp ./Phenotype.txt ./test.gff Si9g49990

For the graphical user interface, ………….

## Output and analyzing a GWAS investigation

The output includes a txt file of haplotypes with detailed information and three pdf files of figures (Fig. 1c-f). The result of haplotypes includes References allele, Alternative allele, Allele Frequency, SNP annotation, SNP positions and haplotypes (Fig. 1d). The information for each haplotype also includes Number of varieties, Varieties ID and its phenotype, Average and SD of phenotype (Fig. 1d).

As an example, we investigated a GWAS result of foxtail millet (Unpublished). The result of this GWAS includes ∼3679 K GWAS SNP *P*-values and 531 SNPs with *P*-value < 9.42 × 10^−7^, and LD is 50 kb. We studied all SNPs that are in the LD 50 kb region of 9:54583294, that is the most significant SNPs (*P*-value = 1.23 × 10^−8^). CandiHap identified one candidate causal gene (*Si9g49990*) (Fig. 1d). SNP 9:54605172 is in LD (50 kb), which is with genome-wide significance in the original GWAS (*P*-value = 1.03 × 10^−7^), and it is a stop gain (Fig. 1d). The boxplot of *Si9g49990* showed significant difference in the phenotype of each haplotype between Hap 1, 2, 6 and Hap 3, 4, 5, 7, 8, 9 (Fig. 1f). The results of other genes in the LD region are not shown because of limited space. User can run the test data at https://github.com/xukaili/CandiHap/tree/master/test_data to check those results.

## Discussion

In order to solve the challenge for GWAS data interpretation, our CandiHap tool is developed to identify candidate causal SNPs and gene(s) from GWAS by integrating linkage disequilibrium (LD) analysis, SNP annotation, haplotype analysis and traits statistics of haplotypes. CandiHap is a flexible and user-friendly toolkit, that provides a rapidly solution form GWAS result to candidate causal gene(s), and it will help researchers to derive candidate causal gene(s) for complex traits study. At the time of writing, there is no tool that performs the same function as CandiHap.

The CandiHap could be widely used for the available GWAS investigations. CandiHap supported Windows, Mac and Linux computers and servers in graphical interface and command line, and applied to any other plant, animal or bacteria species. It should be noted that CandiHap is not intended to be used to predict true causal SNPs and gene(s) since for complex traits. So the outputs of CandiHap are candidate causal SNPs and gene(s). An important application of the CandiHap results is to allow investigators to test ‘a priori’ hypothesis concerning pathways by using candidate causal SNPs as the practical starting point.

Intergenic SNPs are SNPs that are located at least 5 kb up- or downstream of a gene. In general, they are not associated with a gene and not located in a known regulatory region. We set a strict default parameter in CandiHap. The parameter limit mapping SNPs to 2000 bp upstream and 500 bp downstream of gene. The default settings ensure that the result is based on the association signals in gene(s) and with statistical significance. Users may also adjust the parameter in ‘CandiHap.pl’.

In the future, CandiHap will be regularly updated, and extended to fulfill more functions with more user-friendly options.

## Acknowledgments

We thank Yibo Li (Huazhong Agricultural University, China) for fruitful discussion and all our colleagues and friends in the Shanxi Agricultural University who helped us test the tool and provided the valuable suggestions. We thank the anonymous reviewers from the Molecular Breeding journal for improving it through their valuable comments and suggestions. We are grateful to people that interact regularly for the improvements of the system.

## Author information

Xukai Li and Zhiyong Shi contributed equally to this work.

## Additional information

### Publisher’s note

Springer Nature remains neutral with regard to jurisdictional claims in published maps and institutional affiliations.

